# A Whole Blood Molecular Signature for Acute Myocardial Infarction

**DOI:** 10.1101/045013

**Authors:** Evan D Muse, Eric R Kramer, Haiying Wang, Paddy Barrett, Fereshteh Parviz, Mark A Novotny, Roger S Lasken, Timothy A Jatkoe, Glenn Oliveira, Hongfan Peng, Jerry Lu, Marc C Connelly, Kurt Schilling, Chandra Rao, Ali Torkamani, Eric J. Topol

## Abstract

Chest pain is a leading reason patients seek medical evaluation. While assays to detect myocyte death are used to diagnose a heart attack (acute myocardial infarction, AMI), there is no biomarker to indicate an impending cardiac event. Transcriptional patterns present in circulating endothelial cells (CEC) may provide a window into the plaque rupture process and identify a proximal biomarker for AMI. Thus, we aimed to identify a transcriptomic signature of AMI present in whole blood, but derived from CECs. Candidate genes indicative of AMI were nominated from microarray of enriched CEC samples, and then verified for detectability and predictive potential via qPCR in whole blood. This signature was validated in an independent cohort. Our findings suggest that a whole blood CEC-derived molecular signature identifies patients with AMI and sets the framework to potentially identify the earlier stages of an impending cardiac event where conventional biomarkers indicative of myonecrosis remain undetected.

**ABBREVIATIONS:**

AMI: acute myocardial infarction
AUC: area under the curve
CAD: coronary artery disease
CEC: circulating endothelial cells
CMP: circulating microparticles
CVD: cardiovascular disease
GSEA: gene set enrichment analysis
qPCR: quantitative polymerase chain reaction
ROC: receiver operator characteristic

## INTRODUCTION

Despite the significant reduction in the overall burden of cardiovascular disease (CVD) over the past decade, CVD still accounts for a third of all deaths in the United States and worldwide each year ^1,2^. While efforts to identify and reduce risk factors for atherosclerotic heart disease (i.e. hypertension, dyslipidemia, diabetes mellitus, cigarette smoking, inactivity) remain the focus of primary prevention, the inability to accurately and temporally predict acute myocardial infarction (AMI) impairs our ability to further improve patient outcomes ^3^. The current diagnostic evaluation for the presence of coronary artery disease relies on functional testing, which detects flow-limiting coronary stenosis, but it has been known for decades that most lesions underlying AMI are only of mild to moderate luminal narrowings prior to acute plaque rupture and not obstructing coronary blood flow ^4–6^. Accordingly, there is an urgent need for improved diagnostics of the underlying arterial plaque dynamics, fissure and rupture ^7,8^. Increased numbers of circulating endothelial cells (CEC) are known to be present not only in patients with AMI but also with unstable angina – marked by the absence of traditional biomarkers of myonecrosis (troponin, CK-MB) - and may provide a window into the pathophysiologic state preceding an acute atherothrombotic event and the development of myonecrosis ^9,10^.

The transition from stable atherosclerotic disease to a ruptured plaque with acute thrombo-occlusive disease is multifactorial and has been the subject of great study. It is thought to involve a combination of physical (sheer stress, thin fibrous cap vulnerability) and biochemical (proinflammatory, vasoactive) factors ^4^. Prior to plaque rupture most atherosclerotic plaques responsible for acute coronary syndromes are not physiologically significant and there is no current diagnostic modality for accurate identification of unstable plaques ^11^. Differential gene expression patterns of leukocytes have previously been used successfully in the assessment of stable coronary disease ^12–14^. Additionally, microarray-derived gene expression patterns in whole blood and PMBCs of patients presenting with AMI have been studied, and CEC-specific gene expression has been examined in patients with metastatic carcinoma and systemic sclerosis ^15–21^. However, the prior studies in AMI were limited by their size and predictive ability. Here, we focus on CECs as a potential source of gene markers for AMI given their temporal elevation in the peri-plaque rupture process. Elevated numbers of CECs have been implicated by our group and others in the pathophysiology leading to acute myocardial infarction ^9,10,22–25^. In fact, while absent in stable angina, increased CECs have been noted not only in AMI, but also in unstable angina, a condition of plaque instability without elevated biomarkers of myonecrosis (troponins, CK-MB) ^9^. Additionally, CEC elevations during AMI are known to be completely independent of the traditional measurements of troponin and CK-MB ^10^. Thus, our primary motivation for initiating our study of gene expression of CECs is that they may be regarded as a biomarker temporally preceding myonecrosis and a transcriptomic signature derived from these cells, and detectable in whole blood, may provide the key to earlier identification of AMI.

## RESULTS

### Enumeration of circulating endothelial cells in patients with acute myocardial infarction

In this study we first assessed CEC counts in AMI patients (n=28) and healthy control volunteers (n=28). CECs were enriched from whole blood using CD146+ immunomagnetic separation and enumerated using the CellSearch system as previously described.^10^ The median CEC count was elevated in AMI patients with 82.5 cells/mL (range, 4 to 650 CEC/mL) whereas the median for healthy volunteers was 9.5 cells/mL (range, 1 to 80 CEC/mL) (p<0.0001 by Mann-Whitney) (Fig. 1). Receiver operating characteristic (ROC) curve analysis demonstrated an AUC of 0.895 (95% CI 0.810 – 0.980, p<0.0001) for CEC enumeration alone for the discrimination of AMI versus healthy volunteers.

**Figure 1.**
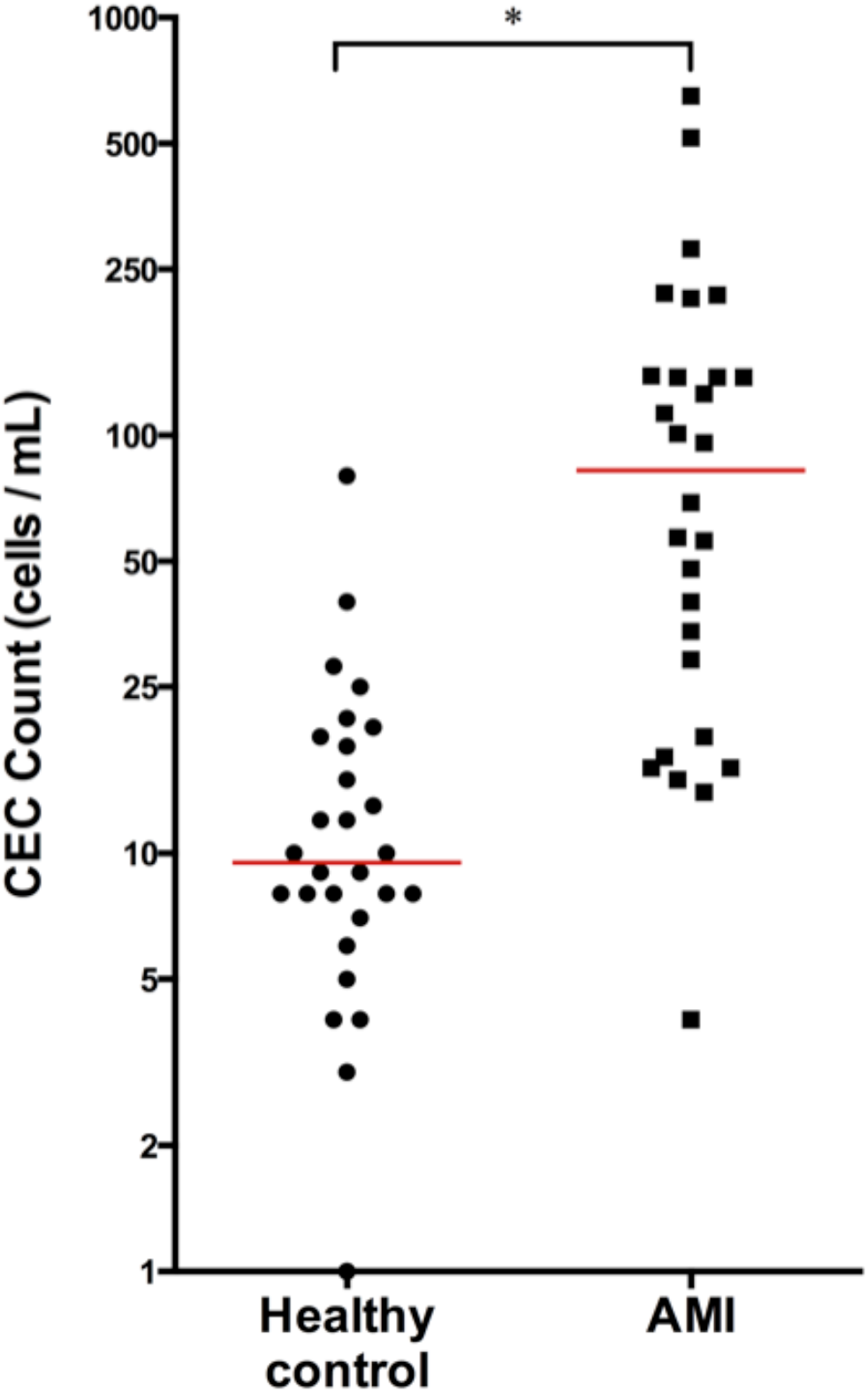
Enumeration of CECs in Patients with AMI. Circulating endothelial cells (CEC) are elevated in the setting of acute myocardial infarction (AMI). CD146+ CECs immuno-magetically separated from whole blood are increased in patients during AMI (n=28) as compared to healthy controls (n=28). * p < 0.0002, non-parametric Mann-Whitney two-tailed t-test.

In support of cellular stress leading to endothelial cell dysfunction and detachment during the acute phase process, we identified circulating microparticles (CMPs), using novel AC electrokinetic methodology previously utilized in the oncology space, as an additional and independent marker for AMI in a separate subset of patients.26 CMPs have previously been shown to be associated with an increased risk of CVD and adverse cardiovascular clinical outcomes in patients with known CAD possibly by promoting procoagulant and inflammatory pathways 27–29. In this group of AMI patients (n=14) and healthy volunteers (n=14), CMPs were elevated in AMI (median 168.5 versus 21.5 particles /mL, p<0.0001 by Mann-Whitney) (**Supplementary Fig. 1A**). Elevated CEC enumeration in AMI was coordinately increased in the same subset of subjects (**Supplementary Fig. 1B**). In these subjects for which both CMP and CEC enumeration was performed, the CMP and CEC counts were highly correlated as measured by Pearson r analysis (R-squared 0.692, p<0.0001) (**Supplementary Fig. 1C**) with no significant differences in their ability to differentiate AMI from control in ROC-curve analysis (AUC for CMP 0.898, 95% CI 0.781 −1.0 AUC for CEC 0.888, 95% CI 0.767-1.0) (**Supplementary Fig. 1D**).

### Microarray gene expression dynamics of enriched circulating endothelial cells

CEC and CMP enumeration is not a practical marker for rapid turnaround in the acute care setting and thus we turned our attention to gene expression assessment. We took an extreme phenotype study design to discover markers in CECs indicative of AMI and detectable in whole blood, and validated their discriminative potential in well-matched subjects. Samples were enriched for CD146+ CECs by the Veridex CellSearch system ^10^ and gene expression determined via microarray. Markers were initially filtered based on biological function (see *Methods)* in order to account for expression differences correlated with co-morbidity differences in our cases vs controls not necessarily indicative of the presence of AMI. Initial marker discovery was performed with elastic net regression in a discovery set of enriched CECs from healthy control volunteers (n=22) and AMI patients (n=21 (Table 1A). The discriminative model trained on this discovery set identified 11 candidate genes (Fig. 2A and Table 2). The top performing marker in the discovery set, heparin-binding EGF-like growth factor (HBEGF), with a coefficient of 0.1132 in our model, was 5.40-fold different in AMI versus controls. However, sulfatase-1 (SULF1) showed the highest fold change, 8.89 (p = 1.97 × 10^−6^), but was less influential on the overall discriminative model (coefficient 0.0283). A model built around the expression levels of these 11 genes effectively discriminated myocardial infarction from healthy control as illustrated in ROC-curve analysis (Fig. 2C).

**Figure 2.**
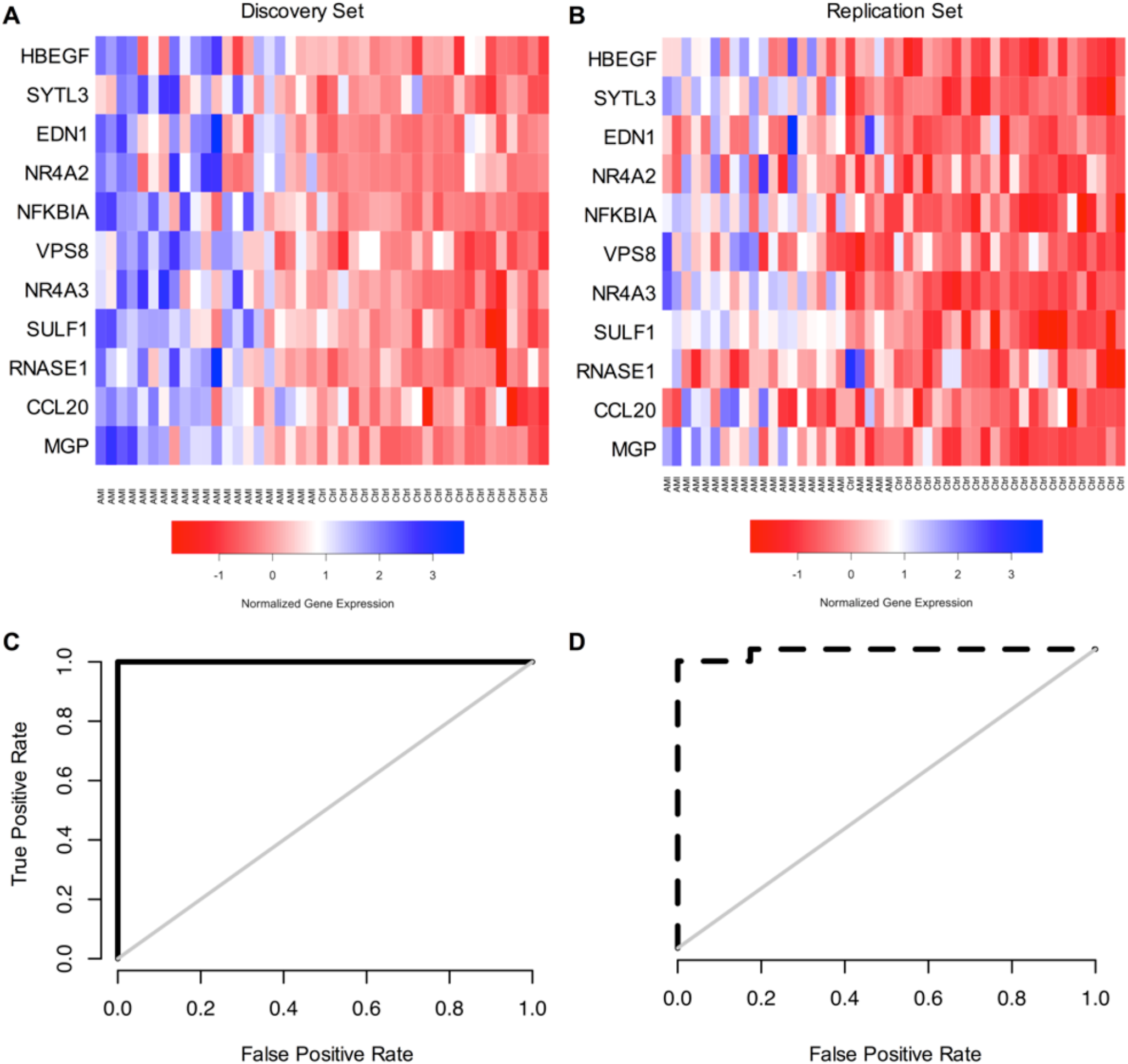
Microarray Analysis of Enriched CECs. An 11-gene signature for AMI was determined from microarray gene expression analysis of enriched CECs from healthy control and AMI patients. (**A, B**) Heat maps for the 11 genes in the microarray of the (**A**) discovery cohort of healthy control (n = 22) and AMI patients (n = 21) and (**B**) replication cohort of healthy control (n = 25) and AMI patients (n = 23) found in the elastic net to discriminate AMI from control. Samples are ordered according to their predicted probability of being an AMI. Expression levels are represented from high (blue) to low (red). (**C, D**) ROC-curves for the 11-gene signature in the (**C**) discovery cohort with AUC of 1.0 (p = 1.90 × 10^−12^ and (**D**) validation cohort with AUC of 0.99 (p = 7.78 x10^−13^).

**Table 1.** Patient Demographics. **(A)**, Age and sex for patients from healthy control and AMI groups used in microarray analysis of enriched CECs. (**B**) Age and sex for patients used in qPCR analysis of whole blood.

**A. Enriched CEC Microarray**
| ***Discovery*** | **Total (n)** | **Male, n (%)** | **Mean Age (years)** |
| --- | --- | --- | --- |
| Control | 22 | 9 (41%) | 28.6 |
| AMI | 21 | 16 (76%) | 59.0 |
| ***Validation*** |  |  |  |
| Control | 25 | 11 (44%) | 28.0 |
| AMI | 23 | 21 (91%) | 62.0 |

**B. Whole Blood qPCR**
| ***Cohort 1*** | **Total (n)** | **Male, n (%)** | **Mean Age (years)** |
| --- | --- | --- | --- |
| Control | 29 | 14 (45%) | 27.9 |
| AMI | 44 | 39 (89%) | 61.5 |
| ***Cohort 2*** |  |  |  |
| Control | 36 | 18 (50%) | 59.9 |
| AMI | 45 | 26 (58%) | 59.9 |

**Table 2.** Candidate Genes from Microarray. Individual genes from enriched CEC microarray used in the 11-gene model to discriminate AMI from control.

| Gene |  | Coefficient | Discovery |  |  | Validation |  |  |
| --- | --- | --- | --- | --- | --- | --- | --- | --- |
|  |  |  | Fold-Change | p-value | adjusted p-value | Fold-Change | p-value | adjusted p-value |
| HBEGF | heparin-binding EGF-like growth factor | 0.1132 | 5.40 | 7.40E-10 | < 0.0005 | 5.16 | 1.6E-06 | < 0.0005 |
| SYTL3 | synaptotagmin-like 3 | 0.0991 | 3.74 | 7.59E-08 | < 0.0005 | 2.17 | 1.1E-02 | 0.088 |
| EDN1 | endothelin 1 | 0.0896 | 3.18 | 1.24E-07 | < 0.0005 | 1.47 | 1.1E-01 | 0.295 |
| NR4A2 | nuclear receptor subfamily 4, group A, member 2 | 0.0583 | 5.80 | 4.24E-08 | < 0.0005 | 11.57 | 2.2E-12 | < 0.0005 |
| NFKBIA | nuclear factor of kappa light polypeptide gene enhancer in B-cells inhibitor, alpha | 0.0563 | 3.55 | 2.05E-07 | < 0.0005 | 5.41 | 1.4E-10 | < 0.0005 |
| VPS8 | vacuolar protein sorting 8 homolog | 0.0555 | 3.08 | 3.14E-07 | < 0.005 | 1.80 | 2.9E-02 | 0.140 |
| NR4A3 | nuclear receptor subfamily 4, group A, member 3 | 0.0461 | 8.39 | 3.36E-08 | < 0.0005 | 6.04 | 1.2E-07 | < 0.0005 |
| SULF1 | sulfatase 1 | 0.0283 | 8.89 | 1.97E-06 | < 0.005 | 2.74 | 1.9E-03 | < 0.05 |
| RNASE1 | ribonuclease, RNase A family, 1 | 0.0119 | 4.45 | 3.29E-06 | < 0.005 | 2.08 | 6.9E-05 | < 0.05 |
| CCL20 | chemokine (C-C motif) ligand 20 | 0.0014 | 6.23 | 3.65E-06 | < 0.005 | 8.45 | 1.8E-10 | < 0.0005 |
| MGP | matrix Gla protein | 0.0013 | 7.86 | 5.16E-06 | < 0.005 | 5.83 | 9.6E-09 | < 0.0005 |

We next replicated this 11-gene model in a separate cohort of control volunteer (n=25) and AMI patient (n=23) samples acquired, processed and sent for microarray analysis independently of our discovery cohorts (Table 1A, Table 2 and Fig. 2B). Mirroring the excellent performance in the initial discovery cohort, the ROC-curve analysis of this independent replication cohort gave an AUC of 0.99 (p = 7.78 × 10^−13^) (Fig. 2D). It should be noted that while the samples used for microarray analysis were enriched in CECs, CD146 is expressed on a subset of cells other than CECs. Additionally, barcode analysis of the gene expression patterns from the enriched CEC microarray reveals evidence for a mixed-cell population based on an elevated number of total genes expressed ^30^ (**Supplementary Fig. 3)**. As a broad assessment of the general gene pathways altered during AMI we also conducted a gene set enrichment analysis (GSEA) on the microarray data (**Supplementary Fig. 4**). As expected, we find that several reactome pathways, such as hemostasis (NES=3.88, p < 1 × 10^−5^, q < 1 × 10^−5^), platelet aggregation (NES= 3.67, p < 1 × 10^−5^, q < 1 × 10^−5^) and GPCR1 ligand signaling (NES=4.60, p < 1 × 10^−5^, q < 1 × 10^−5^), are highly upregulated in AMI.

### A molecular signature for acute myocardial infarction in whole blood

Following the designation of 11 candidate genes on microarray gene expression analysis of enriched CECs as markers for AMI, we asked if the top performing genes in this molecular signature could be assessed directly from whole blood. By examining the whole blood gene expression patterns we would obviate the specialized cell sorting done prior to microarray. To this end, RNA was isolated from whole blood of the same patients (control and AMI) utilized in the microarray replication study (above) with the addition of 14 new AMI patients following RBC lysis from which cDNA was prepared for qPCR analysis (n = 44 AMI and 29 control) (Table 1B). An important distinction is that while CECs had been specifically enriched from patient blood using CellSearch technology for our microarray analysis, here we used only whole blood. The purpose of this experiment was to simply determine whether the gene signature remains detectable and indicative of AMI in this more convenient sample source.

The expression levels for many of the original genes determined in enriched CEC microarray remained significantly elevated in whole blood samples of patients with AMI compared to healthy control volunteers (Fig. 3A). Heparin-binding EGF-like growth factor (HBEGF) showed the highest discriminatory performance between AMI and healthy control patients (AUC 0.97, 95% CI 0.93-1.00, p<0.0001) in whole blood analysis. In terms of expression differences between AMI and healthy control patients, HBEGF was followed by SULF1 (AUC 0.93, 95% CI 0.86-0.99, p<0.0001), NR4A3 (AUC 0.92, 95% CI 0.87-0.98, p<0.0001), NFKBIA (AUC 0.91, 95% CI 0.83-0.97, p<0.0001), and NR4A2 (AUC 0.90, 95% CI 0.83-0.97, p<0.0001). We retrained the elastic net model using the whole blood qPCR values to account for well established differences between microarray vs qPCR based transcriptomic measurements and eliminate those genes that lose discriminative power in whole blood vs enriched CECs. The elastic net regression retained seven discriminative genes (combined AUC 0.997, 95% CI 0.991-1.00) using HBEGF, NR4A3, RNASE1, SYTL3, SULF1, NFKBIA, and NR4A2 (Fig. 3D, solid black line).

**Figure 3.**
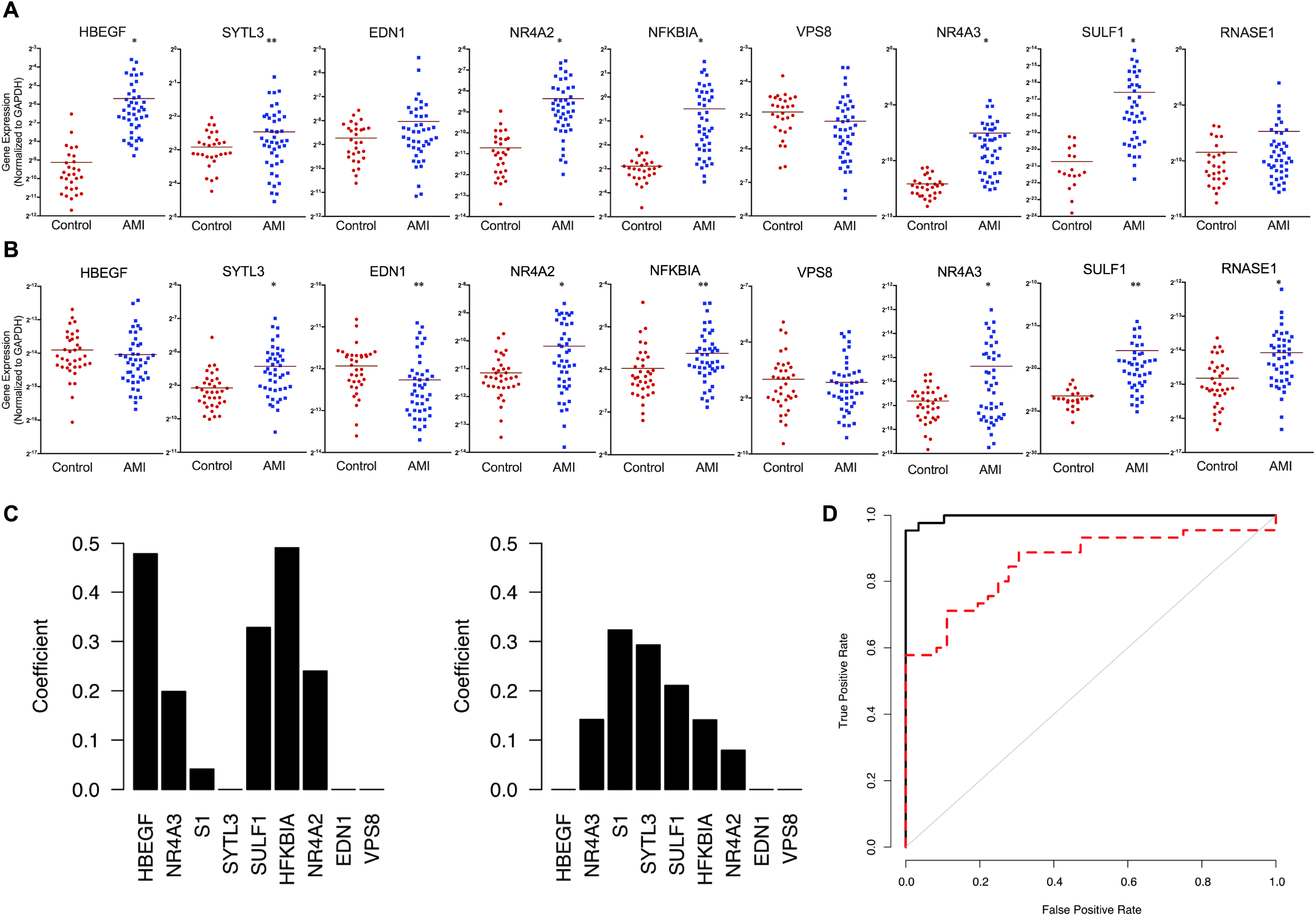
qPCR Analysis of Whole Blood. Candidate genes from enriched CEC microarray were assessed by qPCR in the whole blood of healthy control, stable diseased control, and two separate AMI patient groups. (**A, B**) Individual plots for each gene assessed by qPCR in (**A**) healthy controls (n = 29) vs AMI (n = 44) (cohort 1) and (**B**) diseased controls (n = 36) vs AMI (n = 45) (cohort 2). Specific gene counts normalized by GAPDH for each sample. (**C**) Coefficients for each gene in the model used for cohort 1 and cohort 2. (**D**) ROC-curve analysis for each model: solid black line, trained in cohort 1 and tested in cohort 1; dashed red line, trained in cohort 1 and tested in cohort 2. * p < 0.005, ** p < 0.05, unpaired, two-tailed t-test. Models are evaluated using leave-one-out cross validation when using the same cohort for training and testing.

Finally, given the differences in age, sex and co-morbid diseases apparent in this first cohort of healthy controls compared to AMI patients we validated this gene expression model in a completely independent cohort of patients presenting with AMI (n = 45) as compared to a new cohort of age and sex-matched control patients (n = 36) (Table 1B). The majority of this second control cohort had co-morbid cardiovascular disease with hypertension (n =24, 67%), dyslipidemia (n =27, 75%) and stable coronary artery disease (n = 22, 61%) with many having undergone prior percutaneous coronary intervention (stenting) and/or coronary artery bypass grafting and thus more clinically representative of patients being evaluated for AMI symptoms in an acute care setting (**Supplementary Table 1**). None of the control or AMI patients in this cohort were a part of the cohorts included in microarray studies or the prior qPCR analysis. While the majority of the marker genes performed similarly in this cohort, there were differences, most notably for HBEGF and RNASE1 (Fig. 3B and 3C). Moreover, when we evaluated the gene expression profiles from a subset of cases with reported non-elevated troponins the discriminatory performance of all but these same two genes was modestly improved (**Supplementary Table 2**). The seven-gene discriminative model trained on the original set of AMI patients and healthy control volunteers (cohort one) and validated in this new cohort (cohort two), performed with an AUC of 0.857 (95% CI 0.774-0.941) in ROC-curve analysis (Fig. 3D, dashed red line).

## DISCUSSION

In the acute setting, the diagnosis of AMI relies upon detecting necrotic cardiomyocytes, as reflected by troponin or creatine kinase MB-fraction assays in addition to pathognomonic electrocardiographic changes. Yet each year a number of patients who present to an emergency room with chest pain do not manifest these signs and are discharged, only for some of them to manifest an MI or sudden cardiac death in subsequent days ^31^. Our ultimate goal is to identify a simple, whole blood molecular signature that would not rely upon the endpoint of AMI and myocardial cell death but rather reflect the underlying acute biologic process leading to atherosclerotic plaque rupture and AMI. Here we present the initial steps towards that goal in the designation of a robust gene-based molecular signature for the identification of AMI. We began our search in a specific population of cells, circulating endothelial cells (CEC), that have been identified in increased numbers not only in patients with AMI but also in patients with unstable angina who have not yet manifested biomarker evidence of myonecrosis ^9,32^. As such, CECs can be considered a potential signal of the active peri-plaque rupture process that eventually leads to acute atherothrombotic occlusion of the entire vessel and AMI. While our prior work had validated the findings from Mutin et al. and introduced a novel method for identifying and enumerating CECs, we sought to move beyond enumeration and fully characterize the transcriptome of CECs from patients with AMI so as to generate a specific molecular gene signature that would effectively differentiate AMI from control ^9^. We believe this signature may prove useful for diagnosis of an impending acute coronary syndrome, which will require prospective assessment in at-risk patients who present to an acute care setting with chest pain, suspect AMI, but do not exhibit biomarker signs of myonecrosis.

The initial phase of this study identified 11 genes upregulated in AMI in samples enriched for CECs as determined by gene expression microarray with excellent discrimination. This 11-gene signature was subsequently replicated in an independent cohort of patients with AMI and control volunteers without a loss of power. However, the excellent performance in this initial phase must be tempered by the fact that these comparisons were carried out in patients on separate extremes of the health spectrum: young volunteers without chronic disease and patients presenting with heart attack – a design that may increase statistical power if co-morbidity stratification across the cohorts is appropriately addressed. Additionally, the requirement for specialized cell sorting is a barrier to translating this finding to a point-of-care diagnostic setting.

Accordingly, we then asked if the expression profiles of these genes could be detected from whole blood using qPCR. In whole blood, seven of these genes showed continued expression differences that when analyzed using the elastic net remained significant to the combined molecular signature for discriminating AMI. Further, supporting the non-reliance of this signature on myonecrosis was that the performance of the seven genes of the signature remained unchanged if not marginally superior in a subset of patients presenting to a single center that had no elevation of their cardiac specific biomarkers at the time of presentation.

The determination of candidate genes from microarray analysis was completed by comparing the gene expression dynamics of two very separate populations, healthy controls and patients having AMI. The age and sex differences in addition to the dissimilarities of underlying co-morbid disease or medications of these populations could partly have magnified the discriminative ability of the original 11-gene model in initial testing. The initial AUC values we report in the discovery and validation cohorts in microarray analysis may reflect this magnified discriminative power. However, we would argue that any biases that are not reflective of AMI status would dampen the predictive power observed in our final age and sex matched validation cohort. We addressed this possibility in our final qPCR analysis of the 7-gene model in whole blood using an age and sex matched control cohort of patients with cardiovascular disease for which the model was attenuated though remained significantly robust. Also, given the limited sample size for this study, ethnic differences were not explored.

Currently, there exists no biomarker, diagnostic study or advanced clinical decision making algorithms that foretell a plaque rupture event leading to AMI. Physicians have imperfect tools to calculate ten-year and lifetime risk of potential cardiovascular events based on various epidemiologically derived, population-based risk factors including hypertension, dyslipidemia, diabetes mellitus, age and baseline inflammatory markers, but nothing that places this probability on a more temporal scale ^33^. Additionally, even by using advanced non-invasive imaging tools to identify and then potentially intervening on high risk plaques, those with the greatest potential of rupture or fissure leading to AMI, would not eliminate the majority of future cardiac events ^34^. While gene expression analysis has previously been combined with traditional clinical risk factors to improve determining the likelihood of stable obstructive coronary disease in non-diabetic patients, that classifier does not indicate or predict impending clinical events ^12^. Likewise, several other groups have completed gene expression analysis of whole blood and PBMCs from patients in the setting of AMI to identify the genes with greatest expression differences, but none have reported a similar discriminatory performance as the molecular signature reported here in whether from enriched CEC microarray or whole blood qPCR ^15–18^.

While the inability to accurately identify patients in an acute care setting destined for heart attacks before they fully manifest is a limitation to our study, it is also the driving force behind this study. The logical next step will be the prospective clinical validation of this CEC-derived, whole blood molecular signature for AMI in a large cohort of patients presenting to acute care settings with symptoms and high clinical suspicion for AMI, but without accompanying ECG or biomarker signs of myonecrosis. However, the powerful seven-gene molecular signature presented herein may indeed provide a window into the biologic underpinnings of AMI that may precede current biomarkers and effectively change the way we approach caring for patients with chest pain symptoms in the future.

## MATERIALS AND METHODS

### Patients and control subjects

The study population consisted of patients aged 18-80 years old of both sexes who presented to one of five San Diego County medical centers with the diagnosis of acute myocardial infarction (AMI). Healthy control patients between the ages of 18 and 35 without a history of chronic disease and diseased control patients (with known but stable cardiovascular disease) of between the ages of 18-80 years old were recruited to outpatient clinical centers affiliated with The Scripps Translational Science Institute (STSI) through which Institutional Review Board (IRB) approval for all aspects of this study was obtained. Recruitment of all patients occurred from February 2008 through July 2014. Informed consent was obtained from all subjects in this study. All AMI cases met strict diagnostic criteria including chest pain symptoms with electrocardiographic (ECG) evidence of ST-segment elevation of at least 0.2 mV in two contiguous precordial leads or 0.1 mV in limb leads in addition to angiographic evidence of obstructive CAD in the setting of positive cardiac biomarkers. Our sample sizes were above the calculated threshold of 12-samples at an alpha 0.01, estimated using an established microarray calculator to detect at least two-fold difference with a power of 0.8 and standard deviation of 0.7. This study is registered with ClinicalTrials.gov (NCT01005485).

### Circulating microparticle (CMP) isolation and enumeration

CMPs were isolated from patient plasma using electric current and quantified using a fluorescent microscope with a charge-coupled device camera. Additional details are provided in the Online Appendix.

### Blood collection and CEC sample preparation and enumeration

Arterial blood was collected from AMI patients into both EDTA containing (Becton Dickinson, Franklin Lakes, NJ, USA) and CellSave (Veridex, Raritan, NJ, USA) tubes in the cardiac catheterization laboratory following the placement of an arterial sheath prior to the introduction of any guide wires or coronary catheters. Prior work has shown no effect of access site (venous versus arterial) differences on CEC acquisition ^32^. The samples were maintained at room temperature and processed within 36 hours of collection. The CellTracks®AutoPrep® system was used in conjunction with the CellSearch®CEC kit and the CellSearch®profile kit (Veridex) to immunomagnetically enrich and enumerate CD146+ CECs as previously described ^10,35^. The enriched CEC samples were analyzed with the CellTracks®Analyzer II and the number of CECs in the sample determined. For CEC microarray profiling, the AutoPrep tube with the sample from the CellTracks®AutoPrep® system was removed and placed into the MagCellect Magnet for ten minute incubation. With the tube still in the MagCellect Magnet, the supernatant liquid was aspirated without disrupting the ferrofluid bound cells from which RNA was subsequently isolated. For whole blood samples in EDTA tubes leukocytes and cellular debris was obtained for RNA isolation following RBC lysis with Erythrocyte Lysis Buffer (Qiagen, Valencia, CA).

### Microarray sample preparation

Microarray analysis was performed in three separate experiments each with even numbers of cases and controls to minimize potential batch effects. Enriched CEC-derived RNA was isolated using Trizol Reagent (Life Technologies, Carlsbad, CA). Labeled target antisense RNA (cRNA) and double stranded cDNA using the Ovation™ RNA Amplification System V2 (NuGEN, San Carlos, CA) was prepared from enriched CEC RNA samples. Purified cDNA underwent a two-step fragmentation and labeling process using the Encore Biotin Module (NuGEN). The amplified cDNA targets were hybridized to Affymetrix human U133 Plus 2.0 array to assess expression levels of over 47,000 independent transcripts (Affymetrix, Santa Clara, CA). Following hybridization, arrays were washed and stained before scanning on the Affymetrix GeneChip Scanner from which data was extracted using the Affymetrix Expression Console. Signal intensities from each array were normalized using the robust multichip average expression measure technique.

### Microarray data analysis

Normalized expression values for the microarrays were calculated using RMA normalization ^36^. Quality controls were conducted with the affy and affyQCReport R packages. A Gaussian mixture clustering of the principal components of the expression data detected eight outliers (five AMI and three control), which were discarded (**Supplementary Fig. 2**). We then removed probe sets that are up-regulated in inflammatory diseases. We then used the discovery set to calculate fold changes for each probe set. Probe sets with a fold change less than two-times were removed. We used elastic net regression and the *glmnet* package in R to build a predictive model for acute myocardial infarction using the microarray data ^37^. The model was trained using the discovery set and then predictions were made for the independent replication set. The performance of the model on the discovery and replication sets was evaluated using receiver-operator characteristic curves and the *pROC* package in R ^38^. A differential expression analysis was run on the discovery and replication sets using the *limma* package in R. For each probe set, a linear model was trained to predict acute myocardial infarction. P-values were calculated using an empirical Bayesian method, which were adjusted using the Bonferroni correction ^39^. A gene set enrichment analysis was run on the combined set of discovery and replication samples ^40^. For the GSEA, each probe’s log fold change was used as the ranking statistic, and the GSEA was set to the “classic” mode. All microarray data are available from the Gene Expression Omnibus database (http://www.ncbi.nlm.nih.gov/geo) under accession code GSE66360.

### cDNA synthesis, pre-amplification and qRT-PCR analysis

First-strand cDNA was synthesized from total RNA using High-Capacity cDNA Archive kit (Applied Biosystems, Foster City, CA). The cDNA was pre-amplified using ABI TaqMan PreAmp (Applied Biosystems) and the selected candidate genes were assessed using the qRT-PCR. PCR data of Ct values were exported for further analysis. ΔCts normalized by GAPDH were applied for all data analysis. An elastic net model was trained using the qPCR data to predict acute myocardial infarction ^37^. The performance of the model was evaluated using leave-one-out cross validation and the receiver-operator characteristic curve. Additional methodological details are provided in the Online Appendix.

## ACKNOWLEDGEMENTS

We appreciate the generosity of the research subjects who made this study possible. We could not have completed this work without the help of a talented team of administrators, clinical coordinators and nurses at STSI including Sharen Knowlton, Sarah Topol, Melissa Sasaki, Sharon Haaser and Michelle Miller. Additional thanks to Rajaram Krishnan, David Charlot and Eugene Tu at Biological Dynamics. This research was supported by funding provided by Veridex, Inc. a wholly owned subsidiary of Johnson - Johnson, Inc., Scripps Health and the linked NIH/NCATS Clinical Translational Science Award 5KL2TR001112 and 5UL1TR001114 to Scripps Translational Science Institute.

## AUTHOR CONTRIBUTIONS

E.D.M., P.B., H.W., M.A.N., R.S.L., M.C.C., K.S., C.R., and E.J.T. organized the project and designed the experiments. H.W., F.P., G.O., H.P., and J.L. were responsible for sample processing and performed all experiments. E.D.M., E.R.K., P.B., H.W., and T.A.J. contributed to the data analysis. While all authors contributed to the process, E.D.M., A.T., and E.J.T. directed the writing and editing of the final manuscript.

## CONFLICTS OF INTEREST

E.D.M., E.J.T., H.W., T.A.J. and M.C.C. are listed as co-inventors on a pending patent for the predictive analysis of myocardial infarction. Biologic Dynamics holds a patent for circulating microparticle isolation.

## REFERENCES

1 Go, A. S. et al. Heart Disease and Stroke Statistics-2014 Update: A Report From the American Heart Association. Circulation 129, e28–e292 (2013).

2 WHO. World Health Organization. Global status report on non-communicable diseases. (2014).

3 Stone, G. W. et al. A prospective natural-history study of coronary atherosclerosis. N. Engl. J. Med. 364, 226–35 (2011).

4 Shah, P. K. Molecular mechanisms of plaque instability. Curr. Opin. Lipidol. 18, 492–9 (2007).

5 Body, R. Emergent diagnosis of acute coronary syndromes: today’s challenges and tomorrow’s possibilities. Resuscitation 78, 13–20 (2008).

6 Maddox, T. M. et al. Nonobstructive coronary artery disease and risk of myocardial infarction. JAMA 312, 1754–63 (2014).

7 Storrow, A. B. & Gibler, W. B. Chest pain centers: diagnosis of acute coronary syndromes. Ann. Emerg. Med. 35, 449–61 (2000).

8 Turnipseed, S. D. et al. Frequency of acute coronary syndrome in patients with normal electrocardiogram performed during presence or absence of chest pain. Acad. Emerg. Med. 16, 495–9 (2009).

9 Mutin, M. et al. Direct evidence of endothelial injury in acute myocardial infarction and unstable angina by demonstration of circulating endothelial cells. Blood 93, 2951–8 (1999).

10 Damani, S. et al. Characterization of circulating endothelial cells in acute myocardial infarction. Sci. Transl. Med. 4, 126ra33 (2012).

11 Arbab-Zadeh, Zadeh & Fuster, V. The Myth of the “Vulnerable Plaque”: Transitioning From a Focus on Individual Lesions to Atherosclerotic Disease Burden for Coronary Artery Disease Risk Assessment. J.Am.Coll.Cardiol. 65, 846–855 (2015).

12 Elashoff, M. R. et al. Development of a blood-based gene expression algorithm for assessment of obstructive coronary artery disease in non-diabetic patients. BMC Med. Genomics 4, 26 (2011).

13 Wingrove, J. A. et al. Correlation of peripheral-blood gene expression with the extent of coronary artery stenosis. Circ. Cardiovasc. Genet. 1, 31–38 (2008).

14 Rosenberg, S. et al. Multicenter validation of the diagnostic accuracy of a blood-based gene expression test for assessing obstructive coronary artery disease in nondiabetic patients. Ann. Intern. Med. 153, 425–34 (2010).

15 Głogowska-Ligus,, J. & Dąbek,, J. DNA microarray study of genes differentiating acute myocardial infarction patients from healthy persons. Biomarkers 17, 379–83 (2012).

16 Kim, J. et al. Gene expression profiles associated with acute myocardial infarction and risk of cardiovascular death. Genome Med. 6, 40 (2014).

17 Kiliszek, M. et al. Altered gene expression pattern in peripheral blood mononuclear cells in patients with acute myocardial infarction. PLoS One 7, e50054 (2012).

18 Muller, O. et al. Transcriptional fingerprint of human whole blood at the site of coronary occlusion in acute myocardial infarction. EuroIntervention 7, 458–66 (2011).

19 Suresh, R. et al. Transcriptome from circulating cells suggests dysregulated pathways associated with long-term recurrent events following first-time myocardial infarction. J. Mol. Cell. Cardiol. 74, 13–21 (2014).

20 Tinazzi, E. et al. Gene expression profiling in circulating endothelial cells from systemic sclerosis patients shows an altered control of apoptosis and angiogenesis that is modified by iloprost infusion. Arthritis Res. Ther. 12, R131 (2010).

21 Smirnov, D. A. et al. Global gene expression profiling of circulating endothelial cells in patients with metastatic carcinomas. Cancer Res. 66, 2918–2922 (2006).

22 Li, C. et al. Detection and validation of circulating endothelial cells, a blood-based diagnostic marker of acute myocardial infarction. PLoS One 8, e58478 (2013).

23 Boos, C. J., Balakrishnan, B., Blann,, a D. & Lip, G. Y. H. The relationship of circulating endothelial cells to plasma indices of endothelial damage/dysfunction and apoptosis in acute coronary syndromes: implications for prognosis. J. Thromb. Haemost. 6, 1841–50 (2008).

24 Hladovec, J., Prerovský, I., Stanĕk,, V. & Fabián, J. Circulating endothelial cells in acute myocardial infarction and angina pectoris. Klin. Wochenschr. 56, 1033–6 (1978).

25 Quilici, J. et al. Circulating endothelial cell count as a diagnostic marker for non-ST-elevation acute coronary syndromes. Circulation 110, 1586–91 (2004).

26 Sonnenberg, A. et al. Rapid electrokinetic isolation of cancer-related circulating cell-free DNA directly from blood. Clin. Chem. 60, 500–509 (2014).

27 Amabile, N. & Boulanger, C. M. Circulating microparticle levels in patients with coronary artery disease: a new indicator of vulnerability? Eur. Heart J. 32, 1958–60 (2011).

28 VanWijk, M. J., VanBavel, E., Sturk, A. & Nieuwland, R. Microparticles in cardiovascular diseases. Cardiovasc. Res. 59, 277–87 (2003).

29 Bernal-Mizrachi, Mizrachi et al. High levels of circulating endothelial microparticles in patients with acute coronary syndromes. Am. Heart J. 145, 962–70 (2003).

30 McCall, M. N., Uppal, K., Jaffee, H. A., Zilliox, M. J. & Irizarry, R. A. The Gene Expression Barcode: leveraging public data repositories to begin cataloging the human and murine transcriptomes. Nucleic Acids Res. 39, D1011–5 (2011).

31 Pope, J. H. et al. Missed diagnoses of acute cardiac ischemia in the emergency department. N. Engl. J. Med. 342, 1163–70 (2000).

32 Damani, S. & Topol, E. Author response to comment on “characterization of circulating endothelial cells in acute myocardial infarction”. Sci. Transl. Med. 4, 149lr4 (2012).

33 Goff, D. C. et al. 2013 ACC/AHA guideline on the assessment of cardiovascular risk: A report of the American college of cardiology/American heart association task force on practice guidelines. Circulation 129, 49–76 (2014).

34 Braunwald, E. Progress in the Noninvasive Detection of High-Risk Coronary Plaques. J. Am. Coll. Cardiol. 66, 347–349 (2015).

35 Rowand, J. L. et al. Endothelial cells in peripheral blood of healthy subjects and patients with metastatic carcinomas. Cytometry. A 71, 105–13 (2007).

36 Irizarry, R. A. et al. Exploration, normalization, and summaries of high density oligonucleotide array probe level data. Biostatistics 4, 249–64 (2003).

37 Friedman, J., Hastie, T. & Tibshirani, R. Regularization Paths for Generalized Linear Models via Coordinate Descent. J. Stat. Softw. 33, 1–22 (2010).

38 Robin, X. et al. pROC: an open-source package for R and S+ to analyze and compare ROC curves. BMC Bioinformatics 12, 77 (2011).

39 Smyth, G. in Bioinforma. Comput. Biol. Solut. Using R Bioconductor (Gentleman,, R., Carey, V., Huber, W., Irizarry, R. & Dudoit,, S.) 397–420 (Springer-Verlag, 2005).

40 Subramanian, A. et al. Gene set enrichment analysis: a knowledge-based approach for interpreting genome-wide expression profiles. Proc. Natl. Acad. Sci. U. S. A. 102, 15545–50 (2005).

